# Chemical proteomic approach for in-depth glycosylation profiling of plasma carcinoembryonic antigen in cancer patients

**DOI:** 10.1101/2023.09.22.558933

**Authors:** Jin Chen, Lijun Yang, Chang Li, Luobin Zhang, Weina Gao, Ruilian Xu, Ruijun Tian

## Abstract

Carcinoembryonic antigen (CEA) of human plasma is a biomarker of many cancer diseases, and its N-glycosylation accounts for 60% of molecular mass. It is highly desirable to characterize its glycoforms for providing additional dimension of features to increase its performance in prognosis and diagnosis of cancers. However, to systematically characterize its site-specific glycosylation is challenging due to its low abundance. Here, we developed a highly sensitive strategy for in-depth glycosylation profiling of plasma CEA through chemical proteomics combined with multi-enzymatic digestion. A trifunctional probe was utilized to generate covalent bond of plasma CEA and its antibody upon UV irradiation. As low as 1 ng/mL CEA in plasma could be captured and digested with trypsin and chymotrypsin for intact glycopeptide characterization. Twenty six out of 28 potential N-glycosylation sites were well identified, which were the most comprehensive N-glycosylation site characterization of CEA on intact glycopeptide level as far as we known. Importantly, this strategy was applied to the glycosylation analysis of plasma CEA in cancer patients. Differential site-specific glycoforms of plasma CEA were observed in patients with colorectal carcinomas (CRC) and lung cancer. The distributions of site-specific glycoforms were different as the progression of CRC, and most site-specific glycoforms were overexpressed in stage II of CRC. Overall, we established a highly sensitive chemical proteomic method to profile site-specific glycosylation of plasma CEA, which should generally applicable to other well-established cancer glycoprotein biomarkers for improving their cancer diagnosis and monitoring performance.

**In Brief:** A chemical proteomic approach for glycosylation profiling of proteins was established for glycosylation characterization of plasma CEA with low abundance. Although CEA has been widely used in diagnosis and prognosis of many cancers, it lacks specificity and sensitivity. We found that the glycosylation of CEA on intact glycopeptide level provided additional dimension of molecular features to improve the performance of CEA in cancer diagnosis and progression.

**Highlights:**

- A chemical proteomic approach for glycosylation profiling of proteins with low abundance
- Glycosylation identification of plasma CEA on intact glycopeptide level with high sensitivity and reproducibility
- Glycosylation features of plasma CEA in cancer patients with CRC and lung cancer and in CRC patients at different progression stages

**Graphical Abstract:** 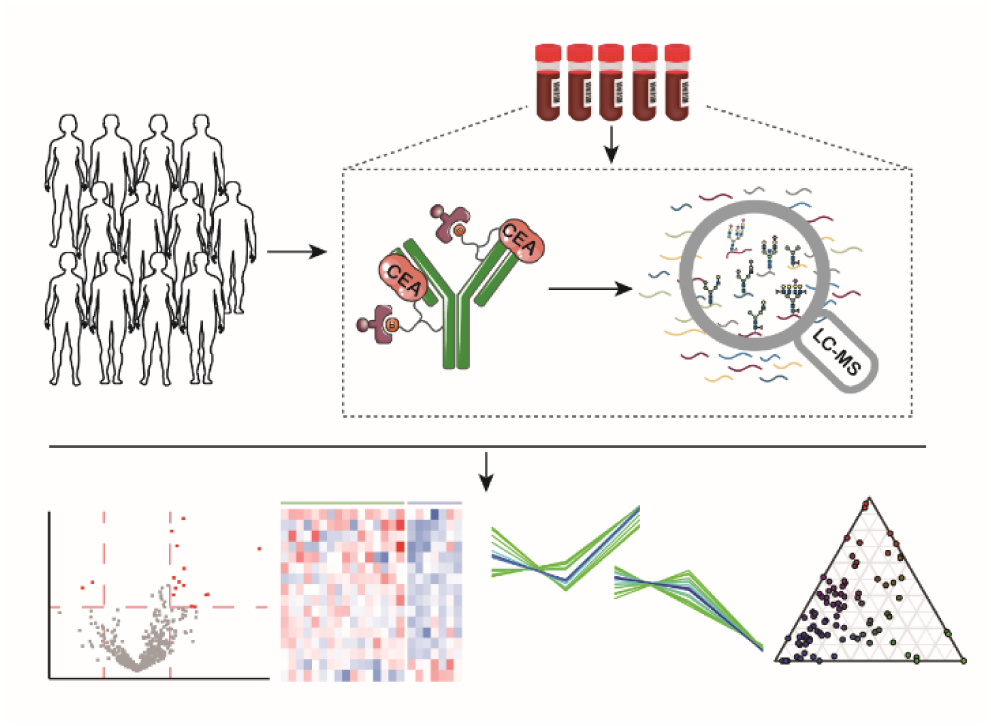

## Introduction

Protein glycosylation is a crucial post-translational modification in health and disease^[1]^. Evidences showed that glycosites and glycan structures are both essential for glycosylation and various biological processes. For example, N-glycosylation regulates cancer metastasis and invasion^[2, 3]^, and the alternation of N-glycan was correlated with the development of cancer^[4]^. PD-L1 contributes to the immune escape of cancer cell and its N-linked glycosylation at site N35, N192, N200 and N219 is crucial for its stability and binding to PD-1^[5, 6]^. Proteins with glycosylation, such as carbohydrate antigens, has been used as cancer biomarkers for decades^[7]^. However, their roles in cancer diagnosis and prognosis are with limited sensitivity and specificity^[8]^. It is highly desirable to provide other molecular features, such as the glycosylation sites and glycans, to increase their performance in cancer diagnosis.

Plasma is the predominant source for clinical diagnostic analyses as its collection is non-invasive, and most FDA-approved plasma biomarkers are with low concentration^[9, 10]^. Functional biomarkers often secrete from tissues into circulation system, but it is challenge to identify them due to their low abundance and the complexity of plasma^[11, 12]^. Antibody is widely used to capture target proteins and applied to immunoprecipitation and ELISA detection, but their affinity is often limited, especially with the presence of complex plasma components^[13, 14]^. Chemical proteomics is a powerful tool for covalently labeling targeted proteins in complex biological systems followed with enrichment and mass spectrometry (MS) analysis^[15]^. For example, an active-site directed probe could be used to profile reactive residues of proteins with high sensitivity^[16]^. We have synthesized a trifunctional probe which could selectively recognize tyrosine phosphorylation (pTyr) and efficiently covalently crosslink pTyr dependent protein complexes in complex clinical samples^[17, 18]^. It is therefore desirable to adopt chemical proteomic approach for profiling targeted proteins and their modifications in plasma samples.

Carcinoembryonic antigen (CEA) serves as a popular biomarker in the diagnosis and prognosis of many cancers such as colorectal cancer (CRC) and lung cancer and its level of > 5 ng/mL was considered abnormal^[19, 20]^. However, the overexpression of CEA could not often be observed in patients with cancer recurrence, and could not apply for early diagnosis of cancer^[21]^. CEA is a highly N-glycosylated protein, which accounts for 60% of molecular mass, and should provide additional dimensional of molecular characteristics for cancer diagnosis and prognosis. Lectin microarray containing 56 plant lectins were used to detect CEA glycosylation in CRC patients, which indicated up-regulation of mannose, N-acetylgalactosamine, N-acetylglucosamine and galactose at stage II of CRC^[22]^. Alternatively, MALDI-TOF-MS has been used to profile 61 unique glycan patterns of purified CEA, which were released by PNGase F^[23]^. To profile the glycosylated sites and glycans simultaneously, intact glycopeptides from 10 µg CEA purified from human CRC were analyzed by capillary electrophoresis-MS system^[24]^. Among 28 potential N-linked glycosylation sites of CEA, 21 of them were successfully identified with the combination of multiple enzymes including trypsin, Glu-C, endoproteinase and pronase^[24]^. Collectively, it is desirable to set up a strategy with high sensitivity for exploring the additional dimension of CEA glycosylation features from clinical samples.

In this study, a chemical proteomic approach was established for in-depth glycosylation profiling of plasma CEA with high sensitivity. Taking advantage of a trifunctional probe, anti-CEA was firstly assembled with the probe which allowed to selectively capture, crosslink and enrich CEA in plasma sample. The intact glycopeptides were released by multi-enzymatic digestion and analyzed by MS (Figure 1). As much as 26 out of 28 N-glycosylation sites were identified in plasma with 500 ng/mL CEA spiked in, and site-specific glycosylation of CEA could be profiled in plasma samples of individual cancer patients with concentration as low as 1 ng/mL. The unique features of site-specific glycoforms of plasma CEA in CRC and lung cancer were successfully characterized. Moreover, its distribution was different in CRC patients at different stage. Therefore, site-specific glycosylation of plasma CEA have the ability to describe the status of disease, and provide potential for cancer diagnosis.

**Figure 1.**
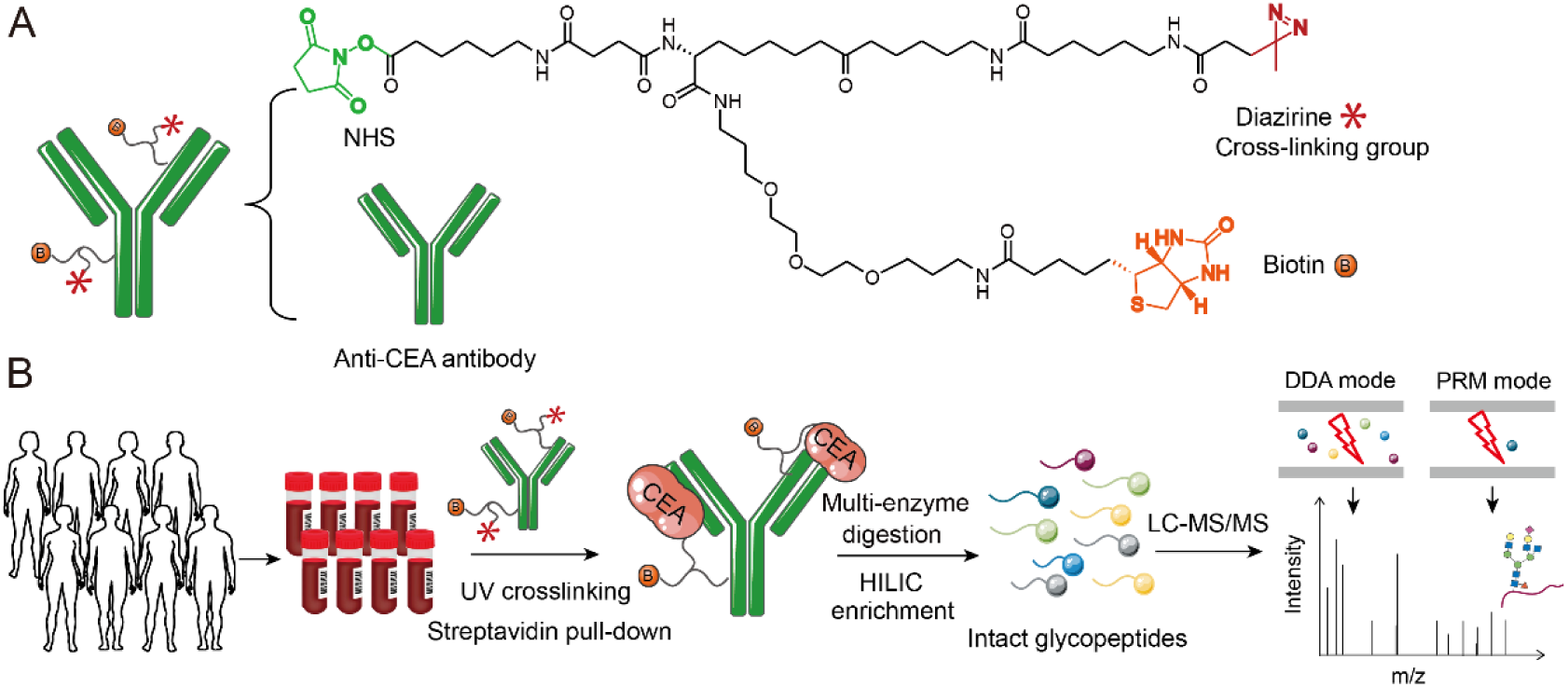
The workflow of this study. (A) The structure of the trifunctional probe which contains NHS, biotin and diazirine group. (B) Plasma was collected and incubated with labeled anti-CEA for capturing plasma CEA, followed with UV cross-linking and multi-enzymatic digestion. Intact glycopeptides were enriched by HILIC and then identified by LC-MS with DDA and PRM mode.

## Experimental Procedures

### Clinical sample

Plasma samples were obtained from the Department of Oncology, Shenzhen People’s Hospital (Guangdong, China). The study was approved by the Institutional Ethical Review Boards of Shenzhen People’s Hospital (number: LL-KY-2021169) and performed according to the Declaration of Helsinki. Plasma samples were collected in EDTA tube at first morning during hospitalization. They were centrifugated within 6 hours, and the upper were divided into two or three copies. They were then stored at - 80°C for further use. Detail information of clinical samples were listed in Table S1.

### Sample preparation

Anti-CEA antibody (#2383, Cell Signaling Technology, the USA) was labeled with the trifunctional probe based on our previous reports with minor modification^[17]^. Briefly, 10 µL antibody was incubated with 1 µL 100 mM probe for 2 min at room temperature and the reaction was stopped by glycine. The labeled anti-CEA was desalted by columns (Zeba^TM^ spin desalting columns, 7K MWCO, #89882, Thermo Scientific^TM^, the USA) and incubated with 1 mL plasma sample with two-fold dilution and 15 µL Sepharose streptavidin beads (#17511301, Cytiva, the USA) overnight. After brief centrifugation, the beads were transferred to 24-well plates for UV irradiation at 365 nm for 10 min at 4 ℃. Next, the beads were washed with washing buffer 1 (RIPA, #P0013B, Beyotime, China) three times, followed with washing buffer 2 (50 mM NH_4_HCO_3_) two times. The captured proteins were alkylated, and digested with trypsin as our previous reports^[25]^. For multi-enzymatic digestion, chymotrypsin (V1061, Promega) or elastase (V1891, Promega) was further added according to the instruction of manufacturer. Finally, the peptides were desalted and intact glycopeptides were enriched by ZIC-HILIC beads (Merck, Product no. 1.50458.0001) according to our previous study^[25]^.

### LC-MS/MS analysis

The intact glycopeptides were analysis by Q Exactive HF-X MS coupled with easy LC system (Thermo Scientific). The nanoLC separation was performed on a 15 cm × 100 μm i.d. C18 (1.9 μm, 120 Å) capillary column at a flow rate of 250 nL/min with a 60 min gradient (5% B to 25% B for 50 min, increase to 90% B at 51 min and hold for 9 min). FA (0.1%, v/v) in water and in 80% ACN (v/v) was used as mobile phase A and B, respectively. All MS spectra were acquired from m/z 400 to 1800 with a mass resolution of 120 000 in DDA mode, in which the 10 most intense ions were selected for MS/MS scan via stepped normalized collision energy (20%, 30% and 40%). Tandem MS was acquired at a resolution of 15 000 and using an isolation window of 2.0 m/z.

Intact glycopeptides from clinical cohort were analyzed by Exploris 240 MS coupled with a Dionex UltiMate 3000 RSLCnano System (Thermo Scientific). The nanoLC separation was performed as following: 5%-28% buffer B for 50 min, 28%– 55% buffer B for 20 min, 55%–99% buffer B for 0.5 min, 99% buffer B for 9.5 min, and 99%-4% buffer B for 0.5min. MS parameters with DDA mode were the same as Q Exactive HF-X MS. In addition, PRM mode was utilized for detecting CEA intact glycopeptides, and the PRM list was shown in Table S2.

### Data processing

The identification of intact glycopeptides were performed by pGlyco3.0^[26]^. The enzymes were set as follows: trypsin, C-term of KR with two missed cleavages; trypsin and chymotrypsin, C-term of KRFYLWM with six missed cleavages; trypsin and elastase, C-term of KRLITSAV with six missed cleavages. Cysteine carbamidomethylation were set as fixed modification, and methionine oxidation and acetylation on protein N-term were set as variable modification. Precursor and fragment tolerance were respectively set as 10 and 20 ppm, and default FDR (1% peptide and glycan) were set. CEA (uniport P06731) was set as database when standard CEA was used and data with PRM mode. Database download from Uniprot-human (in January 2019, 20413 entries) were used when plasma samples were analyzed. Other parameters were set as default.

All the quantification were processed by pGlycoQuant, which could be used for quantitative glycoproteomics at intact glycopeptide level^[27]^. Primary and tandem mass spectrometry were used for intact glycopeptide quantitation. Raw mass spectrometric data and search results from pGlyco3.0 were input and other parameters were set as default. Quantification of site-specific glycoforms could be acquired.

### Experimental design and statistical rationale

In total, plasma from 42 patients were collected and their detail information such as sex, age, cancer type and CEA level were listed in Table S1. Eight out of them with CEA > 500 ng/mL were used for investigating our chemical proteomic approach, and they were measured with three technical replicates. Then, 16 CRC patients and 7 lung cancer patients with CEA > 300 ng/mL were involved to investigate the CEA glycosylation in different cancers, which were identified by LC-MS with DDA mode. Three technical replicates were acquired, and the average intensities of CEA site-specific glycoforms were directly used due to their similar distribution. Site-specific glycoforms were excluded if they were not detected in more than 75% of the samples, and unpaired two-tailed Student’s t-tests were utilized for differential analysis. Finally, 11 CRC patients with different stages, including 3 in stage II (CEA below 2 ng/mL), 3 in stage III (2 with CEA around 3 ng/mL and 1 with CEA 300 ng/mL), and 5 in stage IV (CEA above 250 ng/mL), were investigated. Their CEA glycosylation were detected by LC-MS with PRM mode. No technical replicates were acquired due to biological replicates. Quantified site-specific glycoforms were not further filtered. Volcano, heatmap, Pearson correlation, triangle graph and cluster enrichment were performed using RStudio (RStudio, Boston, MA) with packages, ggplot2, corrplot, ggtern, pheatmap and mfuzz. SIMCA (version 14.1) was used for OPLS-DA analysis.

## Results

### Development of the chemical proteomic approach

We established a chemical proteomic approach for glycosylation of plasma CEA (Figure 1). Firstly, anti-CEA was covalently labeled by the trifunctional probe, which contains NHS, biotin, and diazirine group (Figure 1A). Free primary amines of N terminus or lysine of anti-CEA could be efficiently labeled via N-hydroxysuccinimide ester chemistry, which has been used for capturing pTyr signaling complexes in our previous work^[17]^. Plasma sample of cancer patient was incubated with the labeled anti-CEA for capturing CEA (Figure 1B). Then, covalent bonds between anti-CEA and CEA were efficiently generated via UV cross linking group diazirine with ∼50 Å space arm (Figure 1B). Low-abundant CEA could therefore be enriched from plasma samples by the biotin group. Taking advantage of covalent crosslinking and biotin-streptavidin recognition with high affinity, interferences from plasma sample with extremely high dynamic range could be efficiently removed by stringent washing.

To best explore the molecular signature of site-specific glycosylation, we identified potential glycosylation sites and their glycoforms through multi-enzymatic digestion and ZIC-HILIC-based intact glycopeptides enrichment. Data-dependent acquisition (DDA) couple with liquid chromatography (LC) was firstly applied for identifying and quantifying the enriched intact glycopeptides, and parallel reaction monitoring (PRM) was further adopted for quantified identified intact glycopeptides with improved sensitivity (Figure 1B). Collectively, we designed an integrated chemical proteomic pipeline for efficiently capturing and quantitatively analyzing site-specific glycoforms of CEA.

### Evaluation and optimization of chemical proteomic workflow

Firstly, we investigated the chemical labeling workflow with the readout of identified intact glycopeptides of CEA in plasma and trypsin was used for protein digestion. The number of glycopeptide-spectrum matches (GPSMs) showed nearly two-fold increase, and glycopeptides and site-specific glycoforms exhibited greatly increase with about 70% improvement with UV compared to without UV irradiation (Figure 2A, Table S3). Compare to without UV irradiation, much less peptides from non-specific proteins were observed and GPSMs of CEA accounted for a two-fold higher proportion of all GPSMs with UV (Figure S1). This indicated less peptides from non-specific proteins and glycosylated peptides enrichment greatly improved CEA glycosylation identification. Glycans and glycosites were also greatly increased with UV irradiation. These results indicated that covalent linkage between anti-CEA and CEA greatly contributed to the improvement of glycosylation identification of plasma CEA.

**Figure 2.**
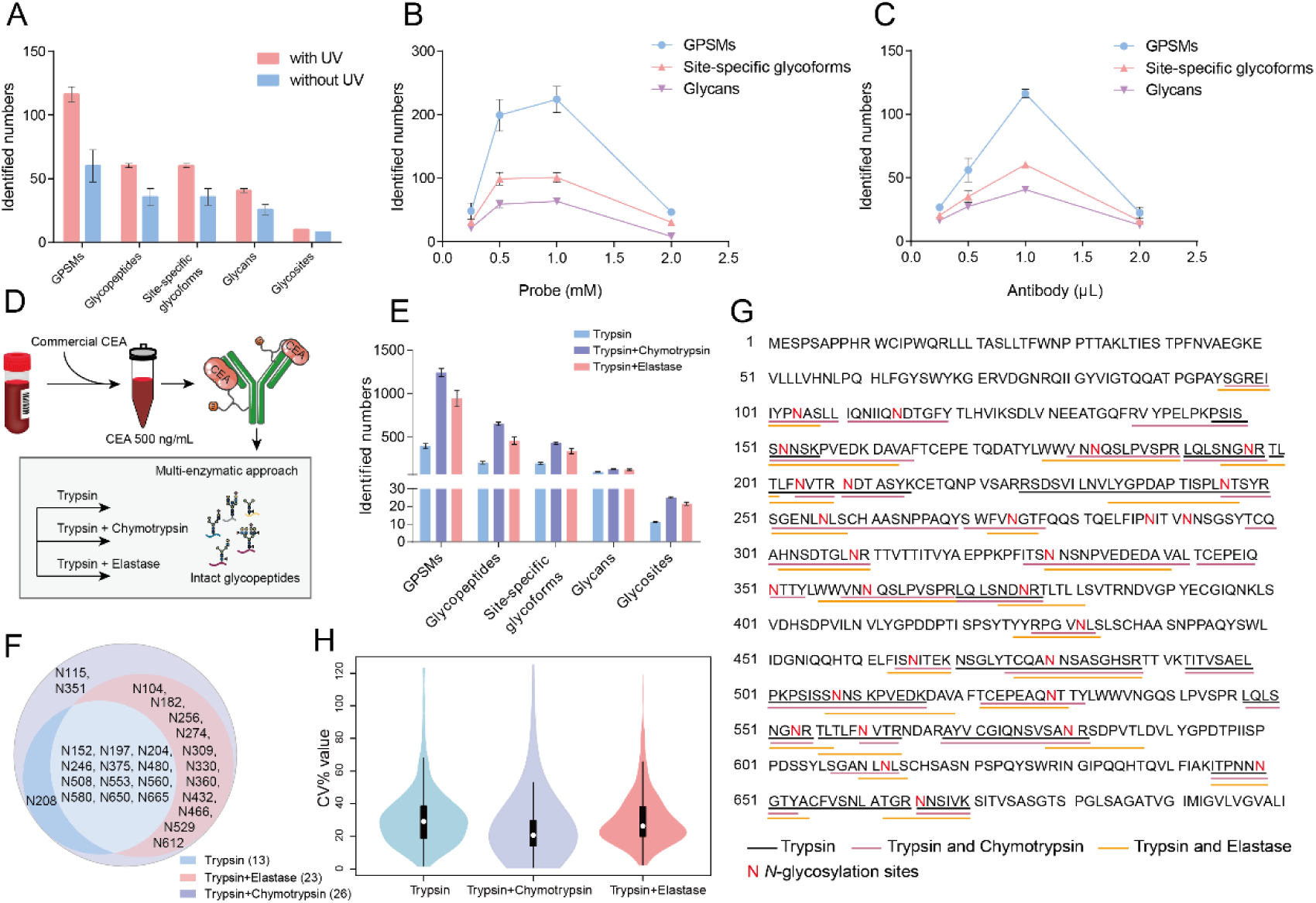
Evaluation and optimization of the workflow for CEA glycosylation identification. The number of glycopeptide-spectrum matches (GPSMs), glycopeptides, site-specific glycoforms, glycans and glycosites were investigated with the condition of chemical labeling (with and without UV) (A), different concentration of probe (B) and antibody (C). Probe (1 mM) and anti-CEA (1 µL) in the reaction with UV crosslinking showed best performance in CEA glycosylation identification. Commercial CEA protein was spiked in plasma with a final concentration 500 ng/mL, followed with multi-enzymatic digestion (trypsin digestion, trypsin and elastase digestion and trypsin and chymotrypsin digestion) for in-depth glycosylation identification (D). Much more GPSMs, glycopeptides, site-specific glycoforms, glycans and glycosites could be identified in trypsin and chymotrypsin digestion (E). In comparison, 13, 23, and 26 CEA glycosylation sites were identified in trypsin digestion, trypsin and elastase digestion and trypsin and chymotrypsin digestion respectively (F). The identified glycosylated peptide sequences with multi-enzymatic approach were shown (G). Three digestion strategies have comparable quantification performance which is acceptable for clinical cohort study (H).

Anti-CEA was labeled with probe via N-hydroxysuccinimide ester chemistry, and its free amino group would be replaced so that its binding affinity might be changed. We thus investigated the labeling reaction and optimized the concentration of probe and antibody for intact glycopeptide identification. The concentration of trifunctional probe greatly affected CEA glycosylation identification. Both 0.5 mM and 1 mM probe had better performance, while 1 mM probe performed the best for identifying slightly more in GPSMs and glycan patterns (Figure 2B, Table S4). The amount of anti-CEA also significantly affected intact glycopeptide identification, among which 1 µL anti-CEA has the best performance for GPSMs, site-specific glycoforms and glycan patterns identification (Figure 2C, Table S5). Therefore, we used 1 mM probe and 1 µL anti-CEA in the reaction for better CEA glycosylation identification.

To increase the depth of CEA glycosylation identification, multi-enzymatic digestion approach was adopted by using trypsin and trypsin paired with either chymotrypsin or elastase. Twenty-six glycosites of commercial CEA were identified in trypsin and chymotrypsin digest as well as trypsin and elastase digest (Table S6), which indicated the in-depth glycosylation coverage of CEA by multi-enzymatic digestion approach. Next, CEA was spiked in plasma with a final concentration 500 ng/mL, and was pulled down by the trifunctional probe-labeled anti-CEA followed with a tryptic digestion and two types of multi-enzymatic digestions (Figure 2D). Compared to trypsin digestion and trypsin combined with elastase digestion, trypsin combined with chymotrypsin digestion showed much more identified GPSMs, glycopeptides, site-specific glycoforms and glycosites (Figure 2E, Table S7). In comparison, 13, 23, and 26 CEA glycosylation sites were identified in trypsin digestion, trypsin and elastase digestion and trypsin and chymotrypsin digestion respectively (Figure 2F), and their identified glycopeptide sequences were shown (Figure 2G). The glycosylation of N288 and N292 could not be observed in our multi-enzymatic approach. Trypsin and chymotrypsin approach has achieved 26 glycosites, which showed the best performance in the glycosylation identification of CEA in plasma as far as we know. Importantly, three digestion strategies have comparable quantification performance which is acceptable for clinical cohort study (Figure 2H, Table S8). Collectively, we adopted trypsin and chymotrypsin for digestion in our workflow for further clinical analysis.

### Profiling of glycosylation of plasma CEA in clinical samples

Our workflow was then applied to 8 individuals whose CEA were above 500 ng/mL and MS with DDA mode was utilized (Table S1, Figure 1B). Expectedly, we identified a large number of GPSMs, glycopeptides, site-specific glycoforms and glycosites in different clinical samples, which indicated the differential glycosylation of plasma CEA between individual patients (Figure 3A, Table S9). Twenty-one CEA glycosylation sites were identified, while N256, N309, N432, N480 and N612 could rarely be observed compared with commercial CEA (Figure 3B). The decrease of identified glycosites might due to the differential CEA glycosylation in individuals. Interestingly, 232 and 169 glycans in plasma and commercial CEA were identified respectively and their quantified CEA levels were roughly equal (Figure S2), demonstrating the much higher diversity of its glycosylation in real world clinical samples. The plasma CEA and the commercial CEA protein did not show great difference in the distribution of modification with oligomannose, complex or hybrid N-glycans. They also displayed nearly the same proportion of fucosylation, which accounted for around 70% of glycan patterns (Figure 3C). Sialylated glycopeptides with fucosylation had nearly 60% increase in plasma CEA compared with commercial CEA, while those without fucosylation occupied similar percentage (Figure 3C). It indicated sialylation peptides of CEA were likely to be modified with fucose in patients with CEA overexpression.

**Figure 3.**
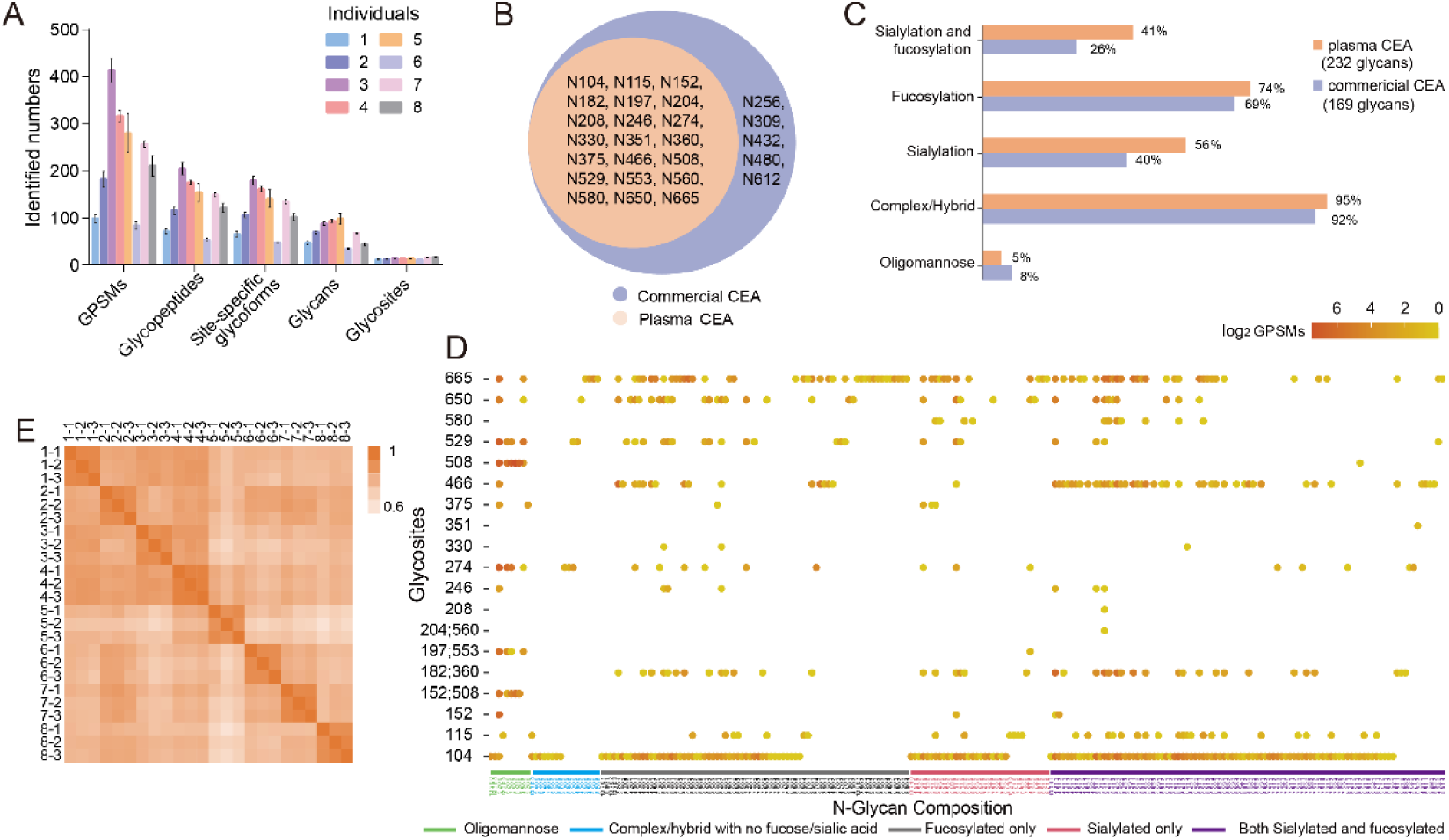
Glycosylation analysis of plasma CEA from 8 individuals. GPSMs, glycopeptides, site-specific glycoforms and glycosites of CEA in different clinical samples were shown in different clinical samples, which indicates their difference (A). Twenty-six and 21 glycosylation sites were identified in commercial CEA and plasma samples respectively (B), and 232 and 169 glycans in plasma and commercial CEA were identified respectively (C). Their identified intact glycopeptides were shown with detailed molecular features. Its color represented their GPSMs (D). The Pearson correlation coefficients of three replicates were high with a range of 0.91-0.97, which indicated high reproducibility (E).

The identified intact glycopeptides were shown with detailed molecular features (Figure 3D). It was observed that N104, N665 and N466 was modified with 168, 85 and 59 types of glycosylation respectively. N508, N197 and N553 were mainly modified with high mannose, while N580 were all modified with sialic acid. It indicated the specificity of glycan composition on sites. We also quantified CEA site-specific glycoforms to evaluate the reproducibility of clinical samples (Table S10). The Pearson correlation coefficients of three replicates were high with a range of 0.91-0.97, which indicated high reproducibility of our method (Figure 3E). However, variations were observed between clinical plasma samples, which showed the heterogeneity between individual patients (Figure 3E). Therefore, our workflow could be used for large-scale analysis of glycosylation of plasma CEA with high reproducibility.

### High-sensitive CEA glycosylation profiling by PRM

Based on the comprehensive discovery of plasma CEA glycosylation, we further evaluated the quantification performance of the workflow which is critical for clinical cohort study with high sensitivity and quantification precision. Patient plasma was diluted according to its CEA concentration of 400, 200 and 100 ng/mL, and its glycosylation were profiled in DDA mode firstly. N529 from the peptide “TCEPEAQNTTY” modified with high mannose Hex(5)HexNAc(2) was identified with excellent MS/MS match when plasma CEA was 100 ng/mL (Figure 4A), and its intensities from chromatographic peaks increased as the increase of plasma CEA (Figure 4B). Additionally, it showed good linearity with R square 0.9976 according to the quantification via pGlycoQuant (Figure 4C). However, only N529 modified with Hex(5)HexNAc(2) and N650 modified with Hex(4)HexNAc(3)NeuAc(1)Fuc(1) could be identified when plasma CEA concentration decreased to 100 ng/mL.

**Figure 4.**
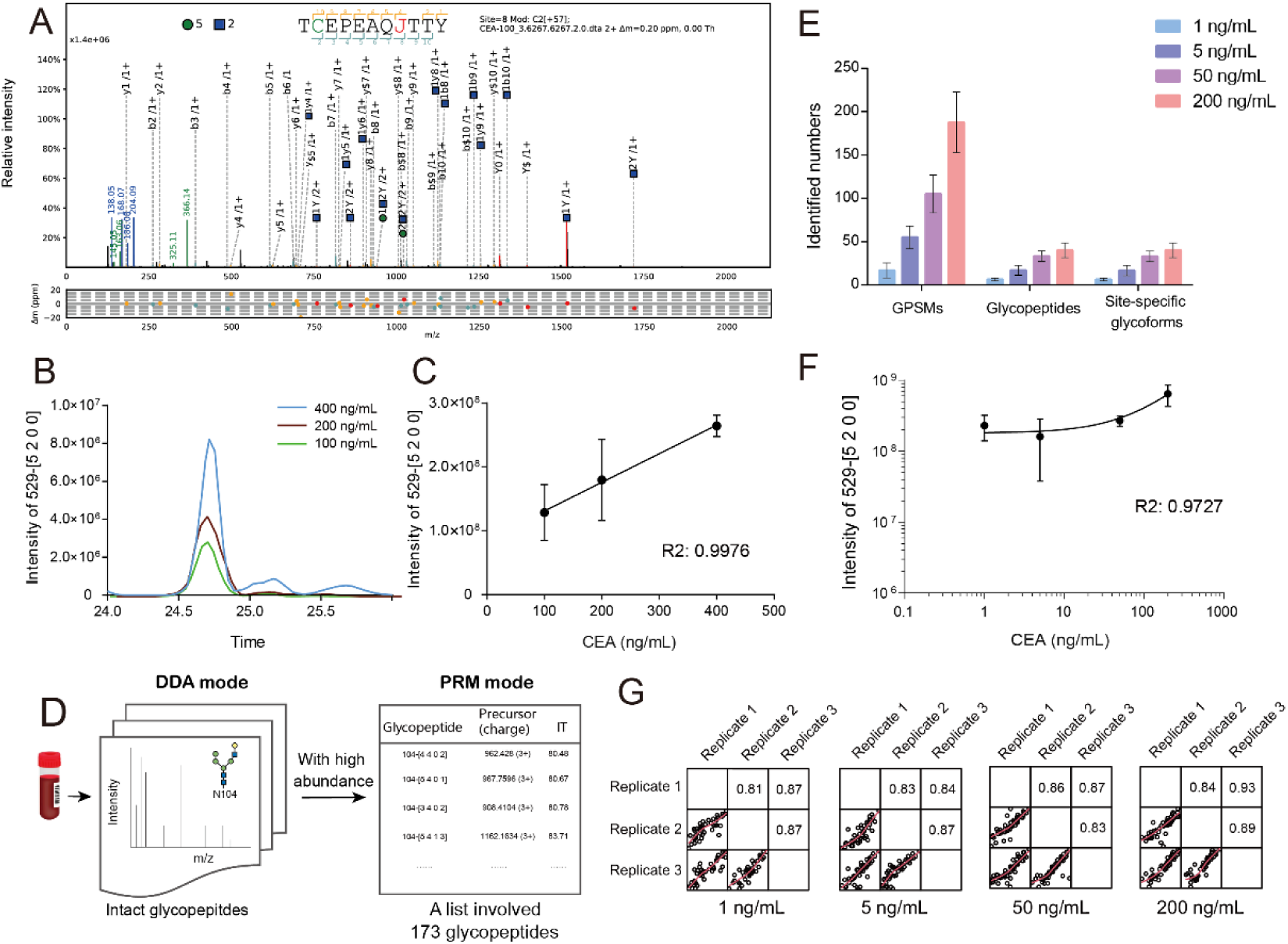
Limit of detection for CEA glycosylation. “TCEPEAQJ(N)TTY” with high mannose Hex(5)HexNAc(2) was identified in DDA mode. Its MS/MS spectral annotation were shown when plasma CEA was 100 ng/mL (A). Its chromatographic peak intensities (B) and linearity (C) showed the ability for quantification. Glycosylation of plasma CEA was then profiled in PRM mode. A list of 173 glycopeptides which have high abundance in plasma CEA identification with DDA mode was generated (D). Their identification was shown when plasma was 1, 5, 50 and 200 ng/mL (E). N529 modified with Hex(5)HexNAc(2) could also be quantified and R square of its linearity was 0.9727 (F). Pearson correlation coefficients for binary comparison of three replicates were calculated when plasma CEA were 1, 5, 50 and 200 ng/mL. Their values were all above 0.8, which indicated the reproducibility of our workflow with PRM mode (G).

To further increase quantification performance and sensitivity, we went on to contract PRM assay for 173 glycosylated peptides (4 are in different charges). These intact glycopeptides were extracted from plasma CEA glycosylation identification above, which have high abundance in DDA mode (Figure 4D, Table S2). The number of GPSMs, intact glycopeptides and site-specific glycoforms reduced as the plasma CEA levels decreased (Figure 4E, Table S11). Thirteen intact glycopeptides could be identified with good precursor peaks and MS/MS matches when plasma CEA was 1 ng/mL (Figure S3), and N529 modified with Hex(5)HexNAc(2) was among them. Its quantification showed good linearity with R square 0.9727 when plasma was 200, 50, 5 and 1 ng/mL (Figure 4F). Additionally, Pearson correlation coefficients of replicates in different CEA concentration were above 0.8, which indicated the reproducibility of the workflow with PRM mode (Figure 4G). Therefore, our chemical proteomic strategy combined with MS analysis in PRM mode could be used for glycosylation profiling of plasma CEA with low concentration.

### Site-specific glycoforms of CEA discriminated patients with CRC and lung cancer

To explore the roles of CEA glycosylation in cancer patients, we investigated their performance in 16 advanced CRC and 7 advanced lung cancer patients with CEA overexpressed (> 300 ng/mL, Table S1), whose age and sex were almost the same (Figure 5A). Intact glycopeptides were detected by LC-MS with DDA mode. Site-specific glycoforms were quantified by pGlycoQuant. Their intensities in 23 patients were draw and showed similar distribution (Figure 5B, Table S12). Based on the quantified site-specific glycoforms, CRC and lung cancer patients could be completely distinguished via OPLS-DA model (Figure S4). According to the criteria that the fold change was > 2 or < 0.5 and T-test showed significance (p<0.05), 16 site-specific glycoforms were observed to be significantly differential from volcano plot (Figure 5C) and their detail intensities were shown (Figure S5). The heatmap described the expressions of significantly differential site-specific glycoforms in the samples, and 14 and 2 site-specific glycoforms were up-regulated in CRC and lung cancer patients respectively (Figure 5D). We depicted the changed sites and plausible glycoforms of CEA (Figure 5E). Twenty-two glycosylation sites of CEA were identified, nine of which (N104, N152, N182, N274, N360, N466, N529, N580 and N665) exhibited great change in different cancers. The glycosylation of N274 and N580 were up-regulated in patients with lung cancer, while others were up-regulated in patients with CRC. Therefore, glycosylation features were correlation with the type of cancers, which might contribute to disease diagnosis.

**Figure 5.**
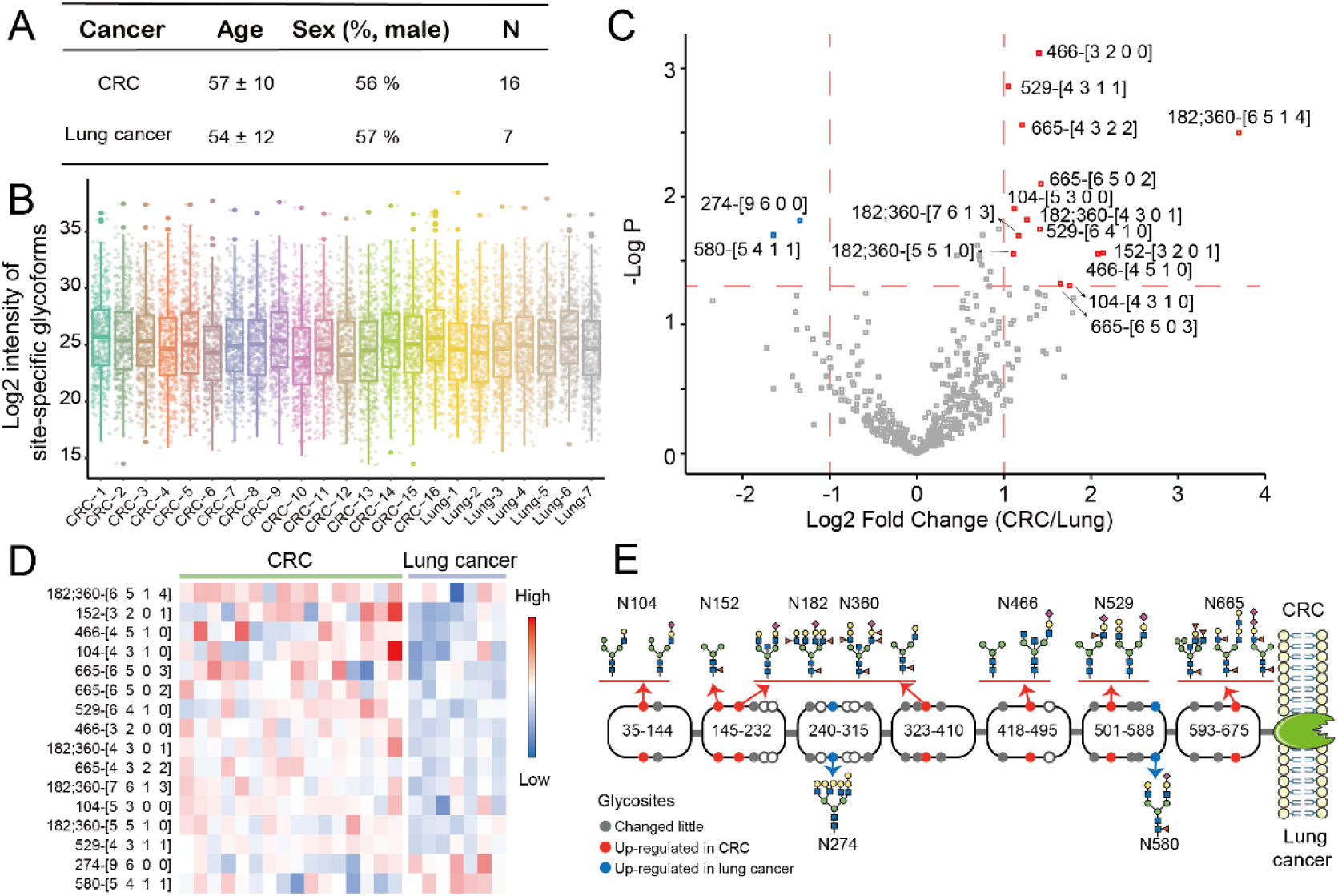
Site-specific glycoforms of plasma CEA in patients with CRC and lung cancer. Patients with 16 advanced CRC and 7 advanced lung cancer were involved (A). Distribution of log_2_-transformed intensities of site-specific glycoforms quantified by pGlycoQuant in 23 patients, and it showed similar distribution (B). According to the criteria that the fold change was > 2 or < 0.5 and T-test showed significance (p<0.05), 16 site-specific glycoforms were observed to be significantly differential from volcano plot. Red and blue refers to up-regulation in CRC and lung cancer respectively (C). Their expressions in different samples were shown in the heatmap (D). The differential sites and plausible glycoforms of CEA were performed. The upper and the lower showed the up-regulation in patients with CRC and lung cancer respectively (E).

### CEA site-specific glycoforms changed as CRC patients in different stages

CEA lacks sensitivity for the diagnosis of CRC, especially in the early stage in which CEA may not be up-regulated. We then investigated the CEA glycosylation of CRC patients with different progression, which contained 3 in stage II (CEA below 2 ng/mL), 3 in stage III (2 with CEA around 3 ng/mL and 1 with CEA 300 ng/mL), and 5 in stage IV (CEA above 250 ng/mL) (Figure 6A, Table S1). We applied LC-MS/MS with PRM mode for its intact glycopeptide analysis. Due to the individual differences, 90 of 173 site-specific glycoforms were identified and quantified (Table S13). Different stages of CRC patients were clustered via OPLS-DA model based on quantified site-specific glycoforms (Figure S6). Also, site-specific glycoforms have differential distribution at different stages (Figure 6B). Although CEA was highly overexpressed in stage IV, most site-specific glycoforms were highly up-regulated in stage II of CRC. It indicated that CEA glycosylation was additional dimension of molecular features to describing the status of diseases in addition to CEA protein level, and we directly used the intensities of CEA glycosylation for analysis. High-mannose glycan on site N466 was expressed only in stage IV, while it on site N208 was not in stage IV. Site-specific glycoforms showed different performances as the progression of CRC, and they were divided into 4 clusters (Figure 6C). Cluster 1 and 2 showed increased trends and dramatically rose at stage IV and III respectively. Cluster 3 and 4 showed decreased trends and greatly dopped at stage III and IV respectively. It was observed that 14, 17, 39 and 20 site-specific glycoforms were respectively enriched in cluster 1, 2, 3 and 4 (Table S14). Detail information of intact glycopeptides in 4 clusters were analyzed (Figure 6D). Most glycosylation of N665 tended to increase in stage III or IV as shown in cluster 1 and 2, while the glycosylation of N650 exhibited decrease in cluster 3 and 4. Most N508 modified with high mannose were enriched in cluster 3 and 4, and N104 modified with sialic acid and fucose were enriched in cluster 3. Most sialylated or fucosylated glycopeptides were enriched in cluster 3, which indicated the sialylation or fucosylation were up-regulated at stage II and greatly declined at stage III. In conclusion, the features of CEA site-specific glycoforms would change as the state of disease, which would distinguish patients and showed the potential for disease diagnosis.

**Figure 6.**
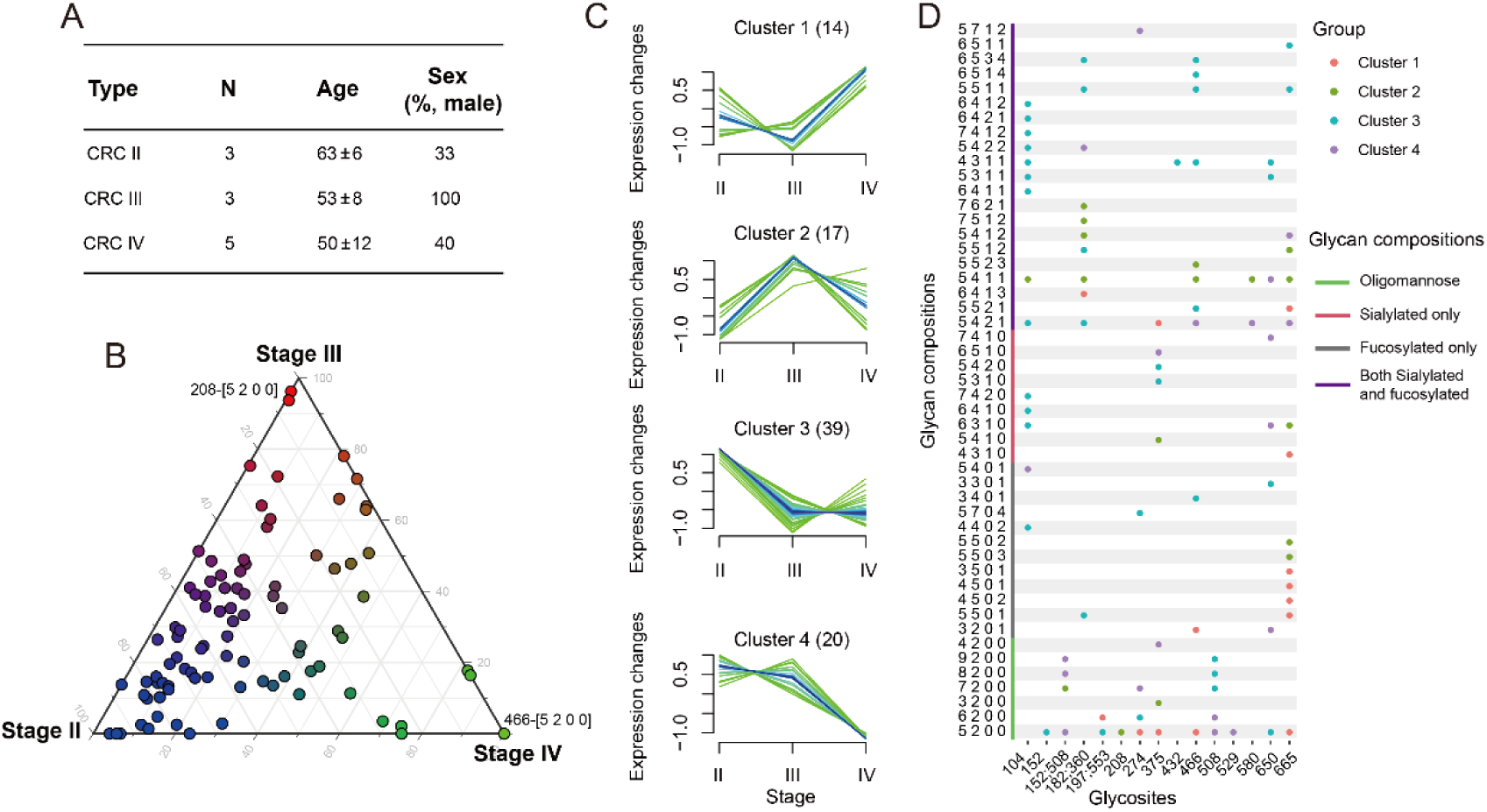
Site-specific glycoforms of plasma CEA in CRC cancer in different stages. CRC patients with different stage were involved, which contained 3 in stage II, 3 in stage III and 5 in stage IV (A). The distribution of site-specific glycoforms of CEA in their plasma were shown (B). Ninety site-specific glycoforms were divided into four clusters, and 14, 17, 39 and 20 site-specific glycoforms were respectively enriched in cluster 1, 2, 3 and 4 (C). The cluster attribution of site-specific glycoforms were shown. The color of dots represented different clusters (D).

## Discussion

In this study, we established a chemical proteomic strategy for glycosylation identification of plasma CEA with high sensitivity. A trifunctional probe has been used to generate a covalent link between anti-CEA antibody and CEA and the strong interaction contributed to the improvement for glycosylation identification of plasma CEA. As much as 26 out of 28 N-link glycosylation sites could be identified and glycosylation of plasma CEA as low as 1 ng/mL could be profiled. The strategy had been applied to 42 clinical samples in this study and the quantification of site-specific glycoforms exhibited high reproducibility, which indicates that it is able to be used for large-scale clinical samples analysis. Our chemical proteomic strategy could be used for glycosylation identification of other biomarkers with low abundance in complex plasma samples.

Multi-enzymatic strategy greatly contributed to in-depth glycosylation identification of plasma CEA. Trypsin combined with chymotrypsin have identified 26 out of 28 N-linked glycosylation sites when FDR was set below 1%, and it showed the most comprehensive N-glycosylation site characterization of CEA on intact glycopeptide level as far as we known. N288 and N292 could not be identified since their MS/MS spectrum were not well matched. They could not either be identified in trypsin, Glu-C or Pronase digestion^[24]^. This may due to the complexity of glycopeptide carried several potential N-glycosylation sites or their absence in the real clinical samples. In addition, some enzymatic glycopeptides would represent two glycosites. Glycopeptide “<u>N</u>VTR” may refer to N204 or N560 due to the multiple cleavage sites, while glycopeptides “SNG<u>N</u>RTLTLF” and “WVN<u>N</u>QSLPVSPRLQL” may be N197 or N553 and N182 or N360, respectively, due to the repetitive regions of CEA. N553 and N560 respectively from glycopeptides “SNG<u>N</u>RTLTLFJVTRNDARAY” and “<u>N</u>VTRNDARAY” could be identified, but it was difficult for us to distinguish N182 and N360.

Sialylation and fucosylation were important features of CEA and they had been shown in 8 individuals who were diagnosed with advanced cancer with metastasis. Sialylation is a biologically important modification, and its alternation during cancer progression has been approved as a potential biomarker^[28]^. Compared to commercial CEA, elevated sialylation of CEA was observed in patients with metastasis. It was consistent with the fact that increased sialylation contributed to metastasis^[29]^. Fucosylation has been involved in the tumor recurrence^[30]^, and core-fucosylation contributed to dismal prognoses and metastatic potential^[31]^. Fucosylation of CEA from 8 individuals was an impressive feature, which accounted for around 70%. Of them, core-fucosylation occupied up to 80%. Terminal fucosylation is involved in the formation of Lewis antigens. CEA carrying sialyl-Lewis x (SLe^x^) is associated with aggressive tumor features and could be a prognosis biomarker^[32]^.

Although CEA has been widely used for cancer diagnosis and prognosis, its glycosylation was overlooked. Our strategy systematically characterized the glycosylation of plasma CEA in cancer patients. The distributions of site-specific glycoforms of plasma CEA differed in patients with CRC and lung cancer as well as in the progression of CRC patients. Most site-specific glycoforms showed up-regulation in CRC patients with stage II, which was consistent with the report illustrating the CEA glycosylation from tumor tissues peaked at stage II during CRC progression^[22]^. Thus, N-glycosylation site characterization of CEA on intact glycopeptide level could be used for evaluation of disease, which could provide additional dimension of features to improve the sensitivity and specificity of cancer diagnosis.

### Conclusions

In conclusion, we develop a chemical proteomic method with high sensitivity and reproducibility to comprehensively explore the glycosylation of plasma CEA based on trifunctional probe and multi-enzymatic approach. Glycosylation of as low as 1 ng/mL CEA could be identified in targeted proteomic analysis mode. This strategy was applied to cancer patients with different stages of CRC and lung cancer. The reliable intact glycopeptide profiling from these clinical samples well support the potential of applying glycosylation features for differentiating cancer patients. Our chemical proteomic strategy therefore provided a generally applicable chemical proteomic method for glycosylation profiling of well-characterized plasma glycoprotein biomarkers.

## Abbreviations

CEA: carcinoembryonic antigen;
DDA: data-dependent acquisition;
PRM: parallel reaction monitoring;
CRC: colorectal cancer;
CV: coefficients of variation;
MS: mass spectrometry;
GPSMs: glycopeptide-spectrum matches;
FDR: false discovery rate;
FA: formic acid;
LC: liquid chromatography;
ACN: acetonitrile;
ZIC-HILIC: zwitterionic hydrophilic interaction liquid chromatography;
HCD: higher energy collision-induced dissociation.

## Acknowledgments

We appreciate the support from the Department of Oncology, Shenzhen People’s Hospital, the Second Affiliated Hospital of Fujian Medical University and Department of Chemistry, Southern University of Science and Technology.

## Author contributions

J. C. conducted data processing and drafted the manuscript. L. Y. implemented the proteomic experiments. C. L. synthesized the trifunctional probe. L. Z. collected the clinical plasma samples. R. X., W. G. and R.T. supervised the study.

